# Long-read genome sequencing and assembly of *Leptopilina boulardi*: a specialist *Drosophila* parasitoid

**DOI:** 10.1101/2020.02.18.953885

**Authors:** Shagufta Khan, Divya Tej Sowpati, Arumugam Srinivasan, Mamilla Soujanya, Rakesh K Mishra

## Abstract

*Leptopilina boulardi* (Hymenoptera: Figitidae) is a specialist parasitoid of *Drosophila*. The *Drosophila*-*Leptopilina* system has emerged as a suitable model for understanding several aspects of host-parasitoid biology. However, a good quality genome of the wasp counterpart was lacking. Here, we report a whole-genome assembly of *L. boulardi* to bring it in the scope of the applied and fundamental research on *Drosophila* parasitoids with access to epigenomics and genome editing tools. The 375Mb draft genome has an N50 of 275Kb with 6315 scaffolds >500bp and encompasses >95% complete BUSCOs. Using a combination of *ab-initio* and RNA-Seq based methods, 25259 protein-coding genes were predicted and 90% (22729) of them could be annotated with at least one function. We demonstrate the quality of the assembled genome by recapitulating the phylogenetic relationship of *L. boulardi* with other Hymenopterans. The key developmental regulators like Hox genes and sex determination genes are well conserved in *L. boulardi*, and so is the basic toolkit for epigenetic regulation. The search for epigenetic regulators has also revealed that *L. boulardi* genome possesses DNMT1 (maintenance DNA methyltransferase), DNMT2 (tRNA methyltransferase) but lacks the *de novo* DNA methyltransferase (DNMT3). Also, the heterochromatin protein 1 family appears to have expanded as compared to other hymenopterans. The draft genome of *L. boulardi* (Lb17) will expedite the research on *Drosophila* parasitoids. This genome resource and early indication of epigenetic aspects in its specialization make it an interesting system to address a variety of questions on host-parasitoid biology.

## INTRODUCTION

Parasitoids are organisms that have a non-mutualistic association with their hosts (Eggleton and Belshaw 1992; Godfray 1994). Around 10%-20% of the described insect species are estimated to be parasitoids. They are spread across five insect orders, i.e., Hymenoptera, Diptera, Coleoptera, Lepidoptera, Neuroptera, Strepsiptera and Trichoptera (Eggleton and Belshaw 1992; LaSalle and Gauld 1991; Heraty 2009), amongst which the vast majority are parasitoid wasps belonging to the order Hymenoptera (LaSalle and Gauld 1991; Godfray 1994; A R Kraaijeveld, Van Alphen, and Godfray 1998). Depending on the stage of the host they attack, they are categorized into the egg, larval, pupal or adult parasitoids. The larvae of parasitoids either feed/develop within the host without impeding its growth (endoparasitic koinobionts) or live on the host after killing or permanently paralyzing it (ectoparasitic idiobionts) (Godfray 1994; A R Kraaijeveld, Van Alphen, and Godfray 1998). Based on the host preference, parasitoids are further classified as generalists and specialists: generalists can parasitize a broad range of species, whereas specialists favor one or two host species (Lee et al. 2009). Likewise, hymenopteran parasitoids display a repertoire of unique features such as polyembryony, hyper-/mutli-/superparasitism, complex multi-level interactions, and haplodiploid sex-determination (Godfray 1994). Many studies have also demonstrated their potential in the biological control of insect pests (Heraty 2009; de Lourdes Corrêa Figueiredo et al. 2015; Smith 1996; Martínez, González, and Dicke 2018; Machtinger et al. 2015; Kruitwagen, Beukeboom, and Wertheim 2018).

*Leptopilina boulardi* (NCBI taxonomy ID: 63433) is a solitary parasitoid wasp from the Figitidae family in the Hymenoptera order (Figure 1). It is a cosmopolitan species, which is ubiquitously found in the Mediterranean and intertropical environments. *L. boulardi* parasitizes *Drosophila melanogaster* and *Drosophila simulans* at second-to early third-instar larval stages and hence, is a specialist (Fleury et al. 2009). However, few strains of the wasp can also infect other Drosophilids like *D. yakuba*, *D. subobscura* and *D. pseudoobscura*, albeit to a lesser extent (Dubuffet et al. 2008; Schlenke et al. 2007). *Leptopilina,* like all the other Hymenopterans, has a haplodiploid sex-determination system. The females are diploid and males are haploid. They are endoparasitic koinobionts, i.e., they lay eggs inside the host larva, allowing the host to grow and feed without rapidly killing it (Fleury et al. 2009; Lee et al. 2009; Kaiser, Couty, and Perez-Maluf 2009). During oviposition, the parasitoid co-inject virulence factors like venom proteins, Virus-like Particles (VLPs) into the larval hemolymph, which helps in taming the host immune responses (Dupas et al. 1996; Goecks et al. 2013; Gueguen et al. 2011). After hatching inside the host hemocoel, the parasitoid larva histolyzes the host tissues gradually. Subsequently, the endoparasitoid transitions into an ectoparasitoid and consumes the host entirely while residing inside the host puparium until emergence. The entire life cycle takes 21-22 days at 25°C (Kaiser, Couty, and Perez-Maluf 2009; Fleury et al. 2009). Alternatively, the host can mount an immune response leading to the death of the parasitoid by encapsulation and the emergence of the host. The interaction, to some degree, also culminates in the death of both host and parasitoid (Fleury et al. 2009; Rizki and Rizki 1990; Small et al. 2012; A R Kraaijeveld, Van Alphen, and Godfray 1998). Such paradigms of evolutionary arms-race are prevalent in insects. However, the combination of *Drosophila* and *Leptopilina*, in particular, has unfolded as a promising tool to study various aspects of the host-parasitoid biology such as coevolutionary dynamics, behavioral ecology, physiology, innate-immune responses, superparasitism (Lee et al. 2009; Fellowes and Godfray 2000; Alex R. Kraaijeveld and Godfray 2009; Tracy Reynolds and Hardy 2004; A R Kraaijeveld, Van Alphen, and Godfray 1998). The advancement could also be attributed to the well-established and extensively studied host.

**Figure 1:**
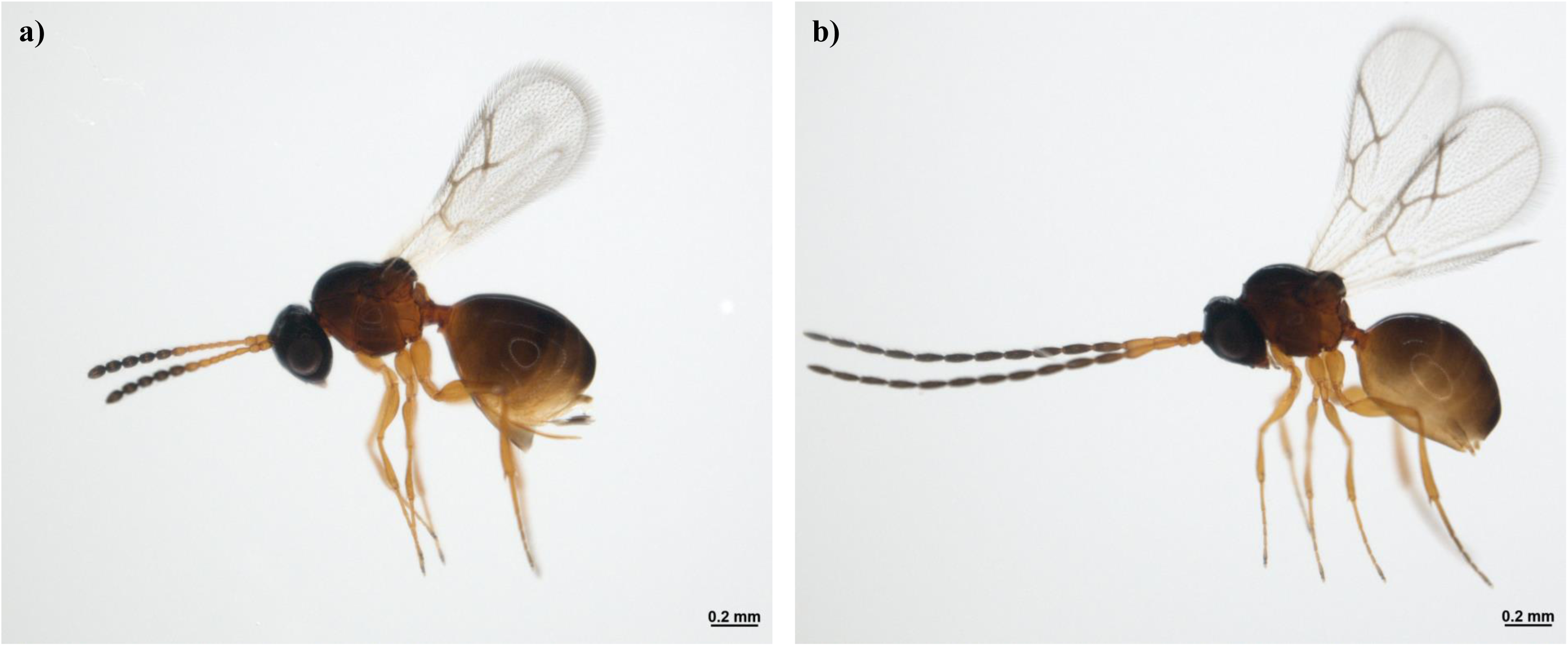
Bright field image of *Leptopilina boulardi* (Lb17 strain). A) Adult female and B) adult male.

The genotype, age, size, and nutritional conditions of the host affect the success of the parasitoid (Boulétreau and Wajnberg 1986; Godfray 1994). Nevertheless, the virulence of the parasitoid is a key factor in determining the fate of an infected host. Although, studies have explored the cause of genetic variance (intra-or inter-specific) and identified the genes involved in the host resistance (A R Kraaijeveld, Van Alphen, and Godfray 1998; Hita et al. 1999; Schlenke et al. 2007; Howell et al. 2012; Salazar-Jaramillo et al. 2017), our understanding of the genetic and epigenetic basis of variation seen in the counter-resistance/virulence of the parasitoids is limited (Alex R. Kraaijeveld and Godfray 2009; Colinet et al. 2010). Another factor that affects the outcome of the host-parasitoid association is the symbiotic partners they harbor, such as *Leptopilina boulardi* Filamentous Virus (LbFV) and *Wolbachia*. LbFV, specific to *L. boulardi*, causes the females to lay eggs in an already parasitized host (superparasitism). Thereby favoring its transmission, and indirectly helping the parasitoid dodge the immune system of the host (Julien Varaldi et al. 2003, 2009; Lepetit et al. 2017; J. Varaldi and Lepetit 2018; Martinez et al. 2012; Patot et al. 2012, 2009). *Wolbachia*, an alpha-proteobacterium, is also the most prevalent endosymbiont of Arthropods. It manipulates the reproductive machinery of the host by inducing either of the following: feminization, male-specific killing, parthenogenesis and cytoplasmic incompatibility, and enhance their transmission to the subsequent generation (Werren, Windsor, and Guo 1995; Vavre, Mouton, and Pannebakker 2009). Hymenopterans, with haplodiploid sex determination, are appropriate hosts for parthenogenesis-inducing *Wolbachia* and have been implicated in the evolution of asexual lineages, such as in *L. clavipes* (K. Kraaijeveld et al. 2016). Interestingly, these bacterial parasites fail to infect the Boulardi clade of the *Leptopilina* genus, unlike the Heterotoma and Clavipes clades (Vavre, Mouton, and Pannebakker 2009; Heath et al. 1999). Such dichotomy observed in the susceptibility of *Drosophila* parasitoids to infections remains elusive.

The epigenetic mechanisms underlying such multispecies interactions that result in the manipulation of behavior and life-history traits of *Leptopilina* genus have not been investigated yet. Therefore, knowing the genomes of *Drosophila* parasitoids will be of great significance in providing a detailed insight into their biology. In this study, we have sequenced the whole genome of *Leptopilina boulardi* (Lb17), generated a high-quality genome assembly, and annotated it. We further looked for its phylogenetic relationship with select metazoans, conservation of genes responsible for body patterning, sex determination and epigenetic regulation of gene expression.

## MATERIALS AND METHODS

### Sample Collection

*L. boulardi* (Lb17 strain), kindly provided by S. Govind (Biology Department, The City College of the City University of New York), was reared on *D. melanogaster* (Canton-S strain) as described earlier (Small et al. 2012). Briefly, 50-60 young flies were allowed to lay eggs for 24 hours at 25°C in vials containing standard yeast/corn-flour/sugar/agar medium. Subsequently, the host larvae were exposed to six to eight male and female wasps, respectively, 48 hours after the initiation of egg lay. The culture conditions were maintained at 25°C and LD 12:12. The wasps (two days old) were collected, flash-frozen in liquid nitrogen, and stored at -80°C until further use.

### Genomic DNA preparation and sequencing

For whole-genome sequencing on Illumina HiSeq 2500 platform (Table 1), the genomic DNA was extracted as follows: 100 mg of wasps were ground into a fine powder in liquid nitrogen and kept for lysis at 55°C in SNET buffer overnight (400 mM NaCl, 1% SDS, 20 mM Tris-HCl pH8.0, 5 mM EDTA pH 8.0 and 2 mg/ml Proteinase K) with gentle rotation at 10 rpm. Next day, after RNase A (100 μg/ml) digestion, Phenol: Chloroform: Isoamyl Alcohol extraction was performed, followed by Ethanol precipitation. The pellet was resuspended in 1X Tris-EDTA buffer (pH 8.0).

**Table 1:** SUMMARY STATISTICS OF THE ASSEMBLED GENOME

| Genome assembly | Numbers |
| --- | --- |
| Assembly size 1n (bp) | 375,731,061 |
| Number of N's (before gapfilling) | 1,423,533 |
| Number of N's (after gapfilling) | 1,216,865 |
| GC content (%) | 28.26 |
| Number of scaffolds | 6315 |
| N50 (bp) | 275,616 |
| Largest scaffold (bp) | 2,405,804 |
| Average scaffold size (bp) | 59,254 |

For long-read sequencing on PacBio Sequel II platform, the genomic DNA preparation was done from 200 mg wasps using the protocol described earlier (Mayjonade et al. 2016) with the following additional steps: Proteinase K digestion for 30 minutes at 50°C after lysis, RNase A digestion for 10-15 minutes at RT (1 μl per 100 μl of 100 mg/ml) after the centrifugation step of contaminant precipitation with potassium acetate and a single round of Phenol:Chloroform:Isoamyl Alcohol (25:24:1, v/v) (Cat. No. 15593031) phase separation before genomic DNA purification using Agencourts AMPure XP beads (Item No. A63880).

### Hybrid genome assembly and assessment of genome completeness

Assembly of the reads was done using a hybrid assembler, MaSuRCA (Zimin et al. 2013). GapFiller (Nadalin, Vezzi, and Policriti 2012) was used to fill N’s in the assembly. Following gap filling, all scaffolds shorter than 500bp were removed from the assembly. The version thus obtained was used for all further analyses. For assessing the quality of the genome assembly, bowtie2 (Langmead and Salzberg 2012) and BUSCOv3 (Simão et al. 2015) was used.

### Identification of repeat elements

To identify repeat elements in the *L. boulardi* assembly, RepeatModeler was used with RepeatScout (Price, Jones, and Pevzner 2005) and TRF (Benson 1999) to generate a custom repeat library. The output of RepeatModeler was provided to RepeatMasker (Tarailo-Graovac and Chen 2009), along with the RepBase library (Bao, Kojima, and Kohany 2015), to search for various repeat elements in the assembly. PERF (Avvaru, Sowpati, and Mishra 2018) was used to identify simple sequence repeats.

### Gene prediction

For RNA-seq based approach, available paired-end data generated from the transcriptome of female *L*. *boulardi* abdomen (SRR559222) (Goecks et al. 2013) was mapped to the assembly using STAR (Dobin et al. 2013). The BAM file containing uniquely-mapped read pairs (72% of total reads) was used to construct high-quality transcripts using Cufflinks (Trapnell et al. 2013). The same BAM file was submitted for RNA-seq based *ab initio* prediction using BRAKER (Hoff et al. 2016). BRAKER uses the RNA-seq data to generate initial gene structures using GeneMark-ET (Lomsadze, Burns, and Borodovsky 2014), and based further uses AUGUSTUS (Stanke et al. 2008) to predict genes on the generated gene structures. In addition to BRAKER, two other *ab initio* prediction tools were used: GlimmerHMM (Majoros, Pertea, and Salzberg 2004) and SNAP (Korf 2004). Using the gene sets generated from various methods, a final non-redundant set of genes was derived using Evidence Modeler (Haas et al. 2008). A protein FASTA file derived using this gene set was further used for functional annotation.

### Gene annotation

BLAST was used to search for homology signatures against SwissProt and TrEMBL databases at an e-value cutoff of 10e-5. InterProScan v5 (Jones et al. 2014) was used to search for the homology of protein sequences against various databases such as Pfam, PROSITE, and Gene3D. The gene ontology terms associated with the proteins were retrieved using the InterPro ID assigned to various proteins.

### Mining of homologs

For protein BLAST (blastp), the proteins were used as query sequences to search against *L. boulardi* proteome. The hit with highest e-value was selected as potential orthologue for a given gene and further subjected to Conserved Domains Search using CDD (Marchler-Bauer et al. 2017) to look for the presence of specific protein domains. Non-redundant BLAST searches at NCBI database were also done to compare with closely associated species from Hymenoptera and other insect orders.

For translated BLAST (tblastn), the proteins were used as query sequences to search against translated *L. boulardi* genome. The hits with e-value greater than 0.01, irrespective of their percentage identity and alignment length, were used for further analysis. The genomic regions that showed matches in tblastn were extended 5 kb upstream and downstream for gene prediction using GENSCAN. The non-redundant peptides obtained from GENSCAN were then subjected to domain prediction using CDD (Marchler-Bauer et al. 2017).

### Multiple sequence alignment and Phylogenetic tree construction

For phylogenetic tree construction of 15 metazoan species, the protein datasets of selected species were downloaded from UniProt (Bateman 2019), choosing the non-redundant proteomes wherever possible. Orthologs were obtained and clustered using OrthoFinder (Emms and Kelly 2015). The tree generated by OrthoFinder was visualized using iTOL v3 (Letunic and Bork 2016). For assigning the putative DNA methyltransferases to DNMT1 and DNMT2 subfamily and aligning the chromodomain/chromoshadow domain sequences obtained by tblastn with seed sequences from *D. melanogaster*, Clustal Omega (Madeira et al. 2019) was used followed by maximum likelihood tree generation with 1000 bootstrap steps using MEGA (Kumar et al. 2018).

### Data availability

The raw reads generated on the Illumina and PacBio platforms are deposited in the Sequence Read Archive (SRA accession SRP144858) of NCBI under the BioProject PRJNA419850. Supplemental material has been uploaded to figshare.

## RESULTS AND DISCUSSION

### Genome assembly and assessment of genome completeness

Previous cytogenetic and karyotypic analysis has estimated the genome size of *L. boulardi* to be around 360Mb (Gokhman et al. 2016). We used JellyFish (Marçais and Kingsford 2011) to determine the genome size of *L*. *boulardi* to be 398Mb. Using the five short-read libraries of ∼200X coverage (70.66GB data) and PacBio reads of ∼30X coverage (10.5GB data) (Supplementary file 1: Table S1), MaSuRCA produced an assembly of 375Mb, made of 6341 scaffolds with an N50 of 275Kb (Table 1). MaSuRCA uses both short Illumina reads and long PacBio reads to generate error-corrected super reads, which are further assembled into contigs. It then uses mate-pair information from short-read libraries to scaffold the contigs. The largest scaffold thus obtained was 2.4Mb long, and 50% of the assembly was covered by 380 largest scaffolds (L50). Using GapFiller, 206Kb out of 1.4Mb of N’s could be filled after ten iterations. From this assembly, all scaffolds shorter than 500bp were removed, leaving a total of 6315 scaffolds.

The quality of the genome assembly was measured using two approaches. First, we aligned the paired-end reads generated from a fresh 250bp library to the assembly using bowtie2 (Langmead and Salzberg 2012). 94.64% of the reads could be mapped back, with 92.32% reads mapped in proper pairs. Next, we used BUSCO v3 (Simão et al. 2015) to look for the number of single-copy orthologs in the assembly. Out of the 978 BUSCOs in the metazoan dataset, 943 (96.5%) complete BUSCOs were detected in the assembly (Table 2). We also performed BUSCO analysis with the Arthropoda (1066 BUSCOs) and Insecta (1658 BUSCOs) datasets and could identify 97% and 95.7% complete BUSCOs in our assembly, respectively (Table 2). The number of complete Insect BUSCOs present in our assembly was similar to that of the other insect genomes (Supplementary file 1: Table S2). Both the results indicate that the generated assembly was nearly complete, with a good representation of the core gene repertoire with only 2.1% and 3.1% of the Arthropod and Insect specific BUSCOs missing from the assembly respectively.

**Table 2:**
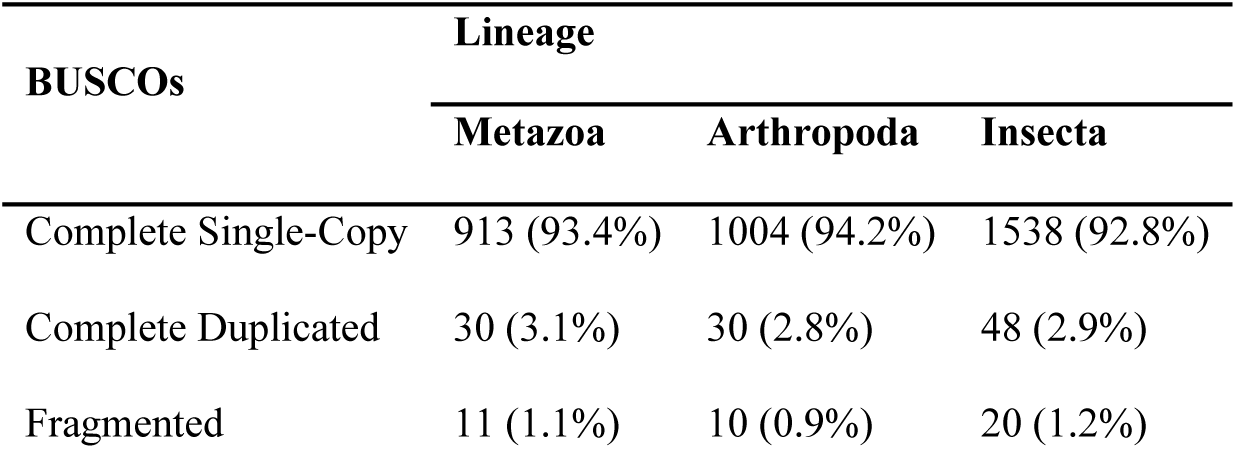
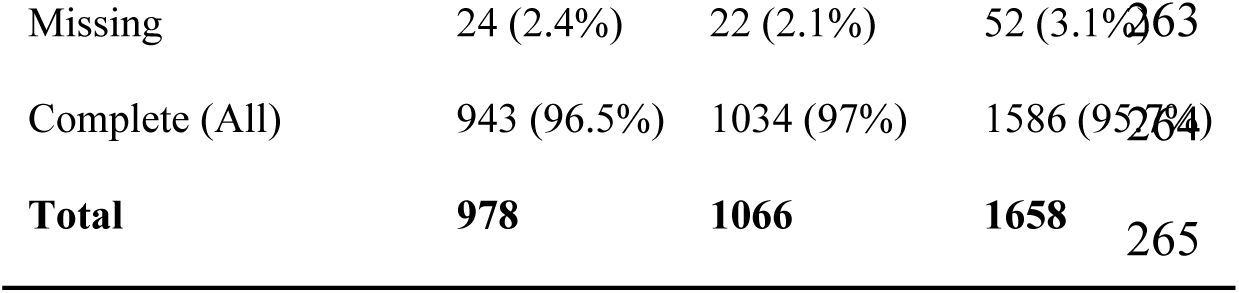
BUSCO ANALYSIS FOR ASSESSING THE COMPLETENESS OF GENOME ASSEMBLY

### Identification of repeat elements

A total of 868105 repeat elements could be identified using RepeatMasker (Tarailo-Graovac and Chen 2009), covering almost 165Mb (43.88%) of the genome. Table 3 summarizes the number of repeat elements identified in the *L. boulardi* assembly as well as their respective types. We further used PERF (Avvaru, Sowpati, and Mishra 2018) to identify simple sequence repeats of >=12bp length. PERF reported a total of 853,624 SSRs covering 12.24Mb (3.26%) of the genome (Table 4). The density of SSRs in the genome of *L. boulardi* was comparable to other insect genomes (Supplementary file 1: Table S3). Hexamers were the most abundant SSRs (40.1%) in the *L. boulardi* genome, followed by pentamers (15.8%) and monomers (14.3%).

**Table 3:** SUMMARY OF REPEAT ELEMENTS IDENTIFIED BY REPEAT MASKER IN THE GENOME

| Repeat Type | Number of Elements | Total Length (bp) | Average Length (bp, rounded) | % Genome Covered |
| --- | --- | --- | --- | --- |
| SINEs | 3721 | 1,651,220 | 444 | 0.44 |
| LINEs | 10573 | 5,613,129 | 531 | 1.49 |
| LTR elements | 12312 | 9,512,954 | 773 | 2.53 |
| DNA elements | 105817 | 31,232,845 | 295 | 8.31 |
| Unclassified interspersed elements | 382214 | 102,924,940 | 269 | 27.39 |
| Small RNA | 186 | 137,204 | 738 | 0.04 |
| Satellites | 2442 | 1,028,732 | 421 | 0.27 |
| Simple repeats | 251669 | 11,461,332 | 46 | 3.05 |
| Low complexity | 46977 | 2,473,942 | 53 | 0.66 |

**Table 4:** DETAILS OF SSRs IDENTIFIED BY PERF IN THE GENOME

|  |  |
| --- | --- |
| Number of SSRs | 853,624 |
| Total Repeat bases | 12.24Mb |
| Repeat bases per Mb genome | 32,587.49 |
| Number of monomers | 122,305 (14.3%) |
| Number of dimers | 101,493 (11.9%) |
| Number of trimers | 72,675 (8.5%) |
| Number of tetramers | 80,493 (9.4%) |
| Number of pentamers | 134,680 (15.8%) |
| Number of hexamers | 341,978 (40.1%) |

### Gene prediction and annotation

Coding regions in the assembled genome of *L. boulardi* were predicted using two different approaches: RNA-seq based prediction and *ab initio* prediction. The number of predicted genes using different method is outlined in Table 5. Using the gene sets generated from various methods, a final non-redundant set of 25259 genes was derived using Evidence Modeler (Haas et al. 2008) (Table 5). The average gene size in the final gene set is ∼3.9Kb. A protein FASTA file was derived using this gene set, which was further used for functional annotation.

**Table 5:** PREDICTION OF GENES IN *L. Boulardi*: SUMMARY OF VARIOUS METHODS

| Evidence Type | Tool | Element | Total Count | Average Length |
| --- | --- | --- | --- | --- |
| **RNA-Seq** | Cufflinks | Gene | 16930 | 10216.46 |
|  |  | Exon | 86962 | 404.44 |
|  | BRAKER | Gene | 45478 | 2461.26 |
|  |  | Exon | 131812 | 384.35 |
| ***ab initio*** | GlimmerHMM | Gene | 28468 | 10529.63 |
|  |  | Exon | 116583 | 243.50 |
|  | SNAP | Gene | 22747 | 856.46 |
|  |  | Exon | 62449 | 222.72 |
| **Combined** | EvidenceModeler | Gene | 25259 | 3886.27 |
|  |  | Exon | 92127 | 333.69 |

The functional annotation of predicted proteins was done using a homology-based approach. 11629 and 19795 proteins could be annotated by performing BLAST against SwissProt and TrEMBL databases, respectively. Further, using InterProScan v5 (Jones et al. 2014), 12,449 out of 25,259 (49.2%) proteins could be annotated with Pfam, while 9346 and 10952 proteins showed a match in PROSITE and Gene3D databases, respectively (Table 6). The gene ontology terms associated with the proteins were retrieved using the InterPro ID assigned to various proteins. A total of 22729 proteins (89.98%) could be annotated using at least one database.

**Table 6:** GENE ANNOTATION OF THE PREDICTED GENES

| Database | Genes Annotated | Percentage Total |
| --- | --- | --- |
| SwissProt | 11629 | 46.04 |
| TrEMBL | 19795 | 78.37 |
| Pfam | 12449 | 49.29 |
| Prosite | 9346 | 37.00 |
| Gene3D | 10952 | 43.36 |
| GO | 9383 | 37.15 |
| Annotated | 22729 | 89.98 |
| **Total** | **25259** | **100.00** |

### Phylogenetic relationship with Hymenopterans

The evolutionary relationship of *L. boulardi* was examined with fifteen metazoan species: one nematode (*C. elegans*), eleven insects – one dipteran (*D. melanogaster*), one lepidopteran (*B. mori*), seven parasitic hymenopterans (*C. solmsi*, *C. floridanum*, *T. pretiosum*, *N. vitripennis*, *M. demolitor* and *O. abietinus*) and two non-parasitic hymenopterans (*P. dominula*, *A. mellifera*), and four chordates (*D. rerio*, *G. gallus*, *M. musculus* and *H. sapiens*) (Supplementary file 1: Table S4). One hundred fifty single-copy orthologs (Supplementary file 2: Figure S1), were obtained and clustered using OrthoFinder (Emms and Kelly 2015), to understand the phylogenetic relationship between the selected species. The tree generated by OrthoFinder was visualized using iTOL v3 (Letunic and Bork 2016). As expected, *L. boulardi* clusters primarily with Hymenopterans and the phylogeny places it as a separate clade and not with other families of Hymenoptera order (Figure 2).

**Figure 2:**
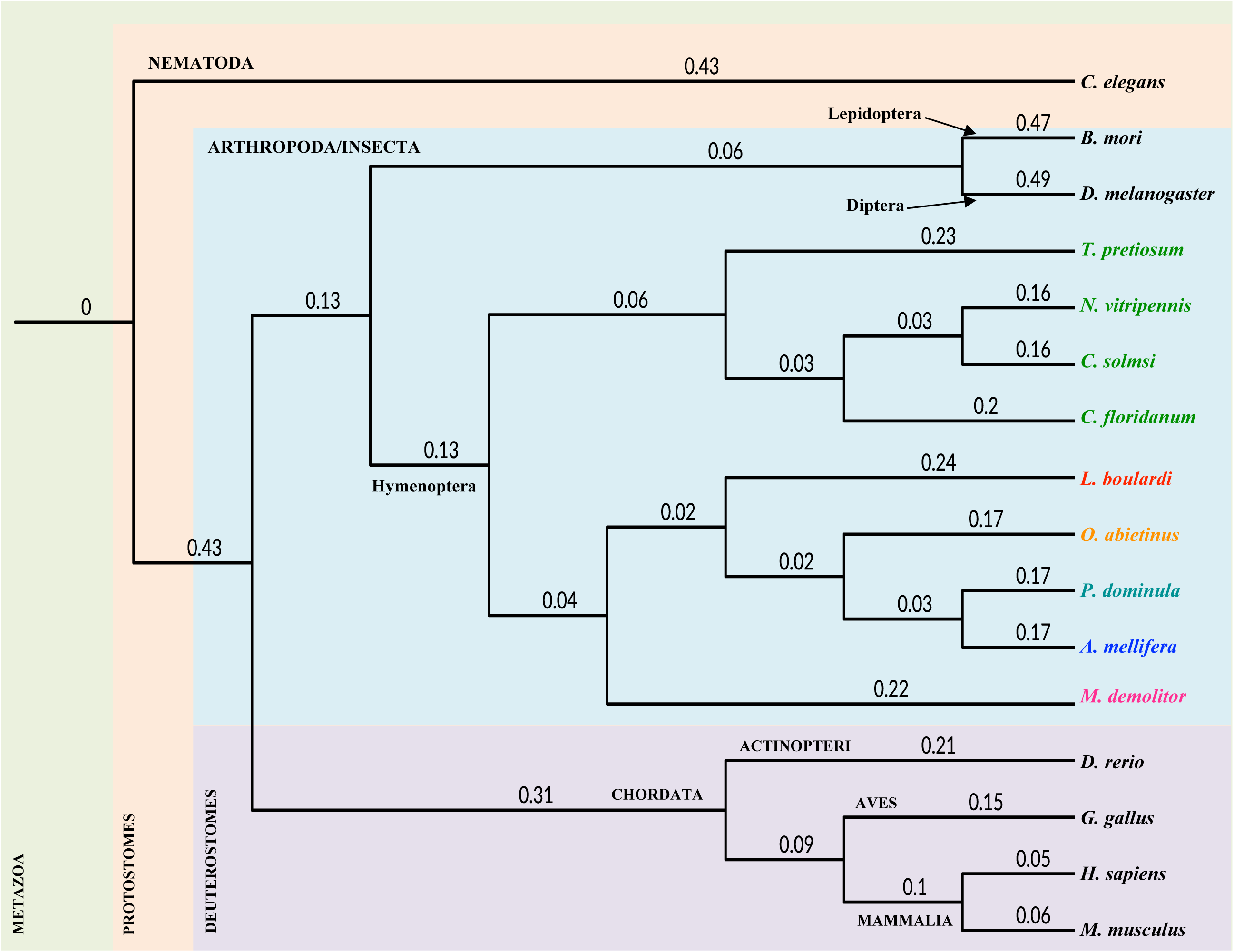
Phylogenetic relationship of *L. boulardi* with selected metazoan species. A phylogenetic tree representing the relationship of *L. boulardi* (red, boldface) with 11 protostomes and four deuterostomes based on 150 single-copy orthologs. Bootstrap values are mentioned at each node. The Phylum/Class is written in uppercase and the order in sentence case. Nine selected species of hymenoptera are shown in different colours based on their superfamily: Orrusoidea (orange), Apoidea (blue), Vespoidea (teal), Chalcidoidea (green), Cynipoidea (red), Ichneumonoidea (magenta).

### Hox genes

Hox genes, a subgroup of Homeobox genes that code for homeodomain-containing transcription factors, play a crucial role during the embryonic development in animals. In addition to their high evolutionary conservation in bilaterian animals, they have received special attention as their genomic arrangement, and expression status determines segment identity along the anterior-posterior body axis (Mallo and Alonso 2013). Unlike vertebrates, where multiple Hox clusters are often found tightly arranged in the genome, the clustering of Hox genes is not very common in invertebrates. The variations observed in the genomic arrangement of Hox genes in insects have helped shed light on the evolution of distinct body plans (Maeda and Karch 2006; Pace, Grbić, and Nagy 2016; Heffer and Pick 2013). Therefore, we investigated the conservation and clustering pattern of Hox genes in the *L. boulardi* genome. The *Drosophila* Hox proteins (Supplementary file 1: Table S4) (Miura, Nozawa, and Nei 2011) were used as query sequences in a protein BLAST to search against *L. boulardi* proteome. All the genes except *Ubx* had full-length protein products in EvidenceModeler gene prediction. For *Ubx*, a full-length protein product was detected in the Cufflinks derived dataset obtained from the available transcriptome of *L*. *boulardi* abdomen (Goecks et al. 2013). In the end, we obtained convincing hits that show high similarity with the Hox proteins of *Drosophila* and Hymenopterans (Supplementary file 1: Table S5 and Supplementary file 3).

The identified Hox genes in *L. boulardi* are spread across four scaffolds. The bithorax complex orthologs – *Ubx*, *abd-A* and *Abd-B* – are located on scaffold00039 (780Kb). However, the orthologs of Antennapedia complex (ANT-C) are distributed in three scaffolds – *pb* and *lab* are located in scaffold00168 (454Kb), *Scr* and *Dfd* are located in scaffold00375 (278Kb), and scaffold00572 (196Kb) contains *Antp*. Overall, the Hox genes are well conserved in *L. boulardi,* span around 1.7Mb of the genome (assuming the scaffolds are contiguous) and are not tightly clustered. To further examine the degree to which Hox genes are dispersed in the genome, the scaffold level draft genome has to be assembled at a chromosome level using techniques such as chromosome linkage mapping, optical mapping, or targeted sequencing of BACs.

### Sex determination genes

Hymenopterans have a haplodiploid sex-determination system wherein the females are diploid, and males are haploid. The diploid females develop from fertilized eggs, whereas the unfertilized eggs give rise to haploid males (arrhenotoky) (Heimpel and de Boer 2008). The two major experimentally supported paradigms of sex determination in Hymenopterans are complementary sex determination (CSD) (Beye et al. 2003) and genome imprinting (Dobson and Tanouye 1998). It has been reported in previous studies that *Leptopilina* genus lacks CSD (Hey and Gargiulo 1985; Biémont and Bouletreau 1980; Van Wilgenburg, Driessen, and Beukeboom 2006) but whether the primary signal for sex determination cascade is the differential methylation status of the maternal and paternal chromosome, is still unclear.

We took the previously described sex determination proteins downstream in the cascade from *D. melanogaster* and *L. clavipes* (Geuverink et al. 2018) and searched for their homologs in *L. boulardi* using blastp approach. We found putative orthologs of the major effector genes (*doublesex and fruitless*) and the genes regulating their sex-specific splicing (*transformer* and *transformer-2*) (Supplementary file 1: Table S6), implying that the downstream cascade of sex determination is well preserved. However, we could only identify one *transformer* gene as opposed to the presence of *transformer* and its paralogue t*ransformerB* in *L. clavipes*.

### DNA methyltransferases

Two families of DNA methyltransferases (DNMTs) are well-known to be responsible for DNA methylation, which occurs primarily at CpG sites in mammals. DNMT3 is a *de novo* methyltransferase, while DNMT1 is known to be involved in the maintenance of DNA methylation (Goll and Bestor 2005). DNMT2, on the other hand, the most conserved methyltransferase in eukaryotes, was initially assigned as a member of DNMT family but later renamed as TRDMT1 (tRNA aspartic acid methyltransferase 1) that justifies its negligible contribution to the DNA methylome (Jurkowski et al. 2008). Other than the role in caste development in social insects (Chittka, Wurm, and Chittka 2012; Herb et al. 2012; Bonasio et al. 2012), cytosine methylation (genome imprinting) has been shown to be the primary signal of the sex determination cascade in the haplodiploid hymenopterans lacking complementary sex determination system (Dobson and Tanouye 1998). In order to assess the methylation status of *L. boulardi* genome, we looked at the CpG content in all exons. Typically, exons that undergo methylation in the genome display an underrepresentation of CpG content due to spontaneous deamination of methylated cytosines into thymines. Hence, genomes that have DNA methylation show a bimodal distribution of CpG content (Elango and Yi 2008), as shown for human and honey bee exons in Supplementary file 2: Figure S2A and B. We observed no such bimodality for exons of *L. boulardi* (Supplementary file 2: Figure S2C).

We further searched for the presence of DNA methyltransferases in *L. boulardi*. Corresponding sequences from *N. vitripennis* (Supplementary file 1: Table S7) were used as seed sequences for identification of DNMTs in *L. boulardi* using blastp and tblastn. We obtained two putative DNA methyltransferases, which were then aligned to DNMTs from *A. mellifera*, *Bombyx mori*, *D. melanogaster*, *N. vitripennis* and *T. pretiosum* (Supplementary file 1: Table S6) using Clustal Omega (Madeira et al. 2019). The maximum likelihood tree thus generated with 1000 bootstrap steps using MEGA (Kumar et al. 2018) assigned the two putative DNA methyltransferases to DNMT1 and DNMT2 subfamily (Figure 3, and Supplementary file 4).

**Figure 3:**
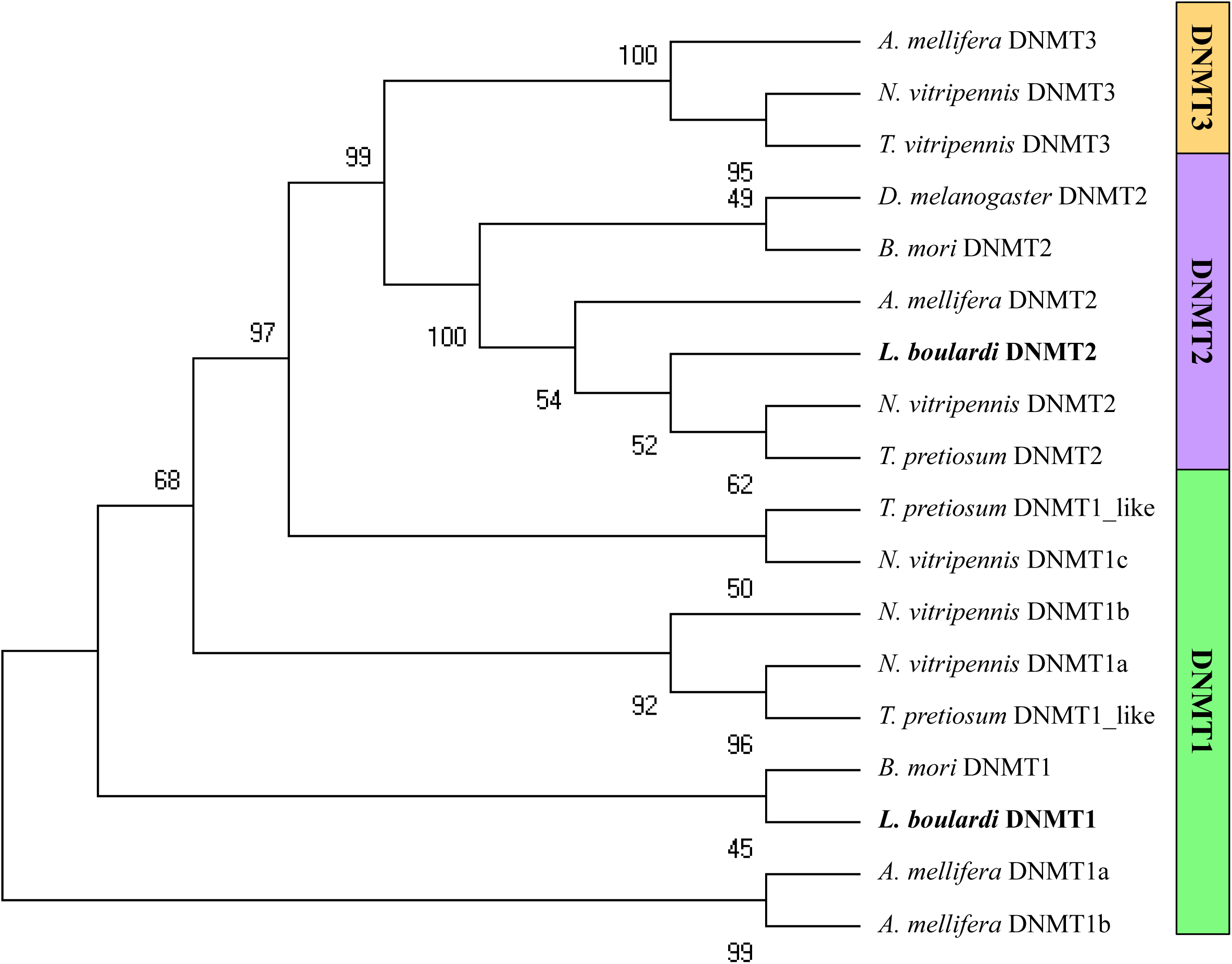
Phylogram of putative DNA methyltransferases in *Leptopilina boulardi*. *L. boulardi* is written in boldface. Bootstrap values are shown at each node. Putative DNMT2 and DNMT1 of *L. boulardi* clusters with the DNMT2 and DNMT1 of other insects, respectively.

An incomplete set of DNA methylation toolkit has been previously described in insects, and it does not always mean an absence of DNA methylation (Glastad, Hunt, and Goodisman 2014; Bewick et al. 2016). The unimodal distribution of observed/expected CpG content in the exons and the presence DNMT1 hints towards possible DNA methylation in non-CpG context in the genome of *L. boulardi*. However, the presence of detectable levels of DNA methylation and the methylation pattern in this genome of *L. boulardi* at different developmental stages needs to be further investigated experimentally.

### Polycomb group, Trithorax group and heterochromatin factors

The expression of genes in the eukaryotes is regulated by numerous evolutionary conserved factors that act in a complex to either direct post-translational modifications of histones or remodel chromatin in an ATP-dependent manner (Grimaud, Nègre, and Cavalli 2006). It is well established that the transcriptionally active states of chromatin are maintained by Trithorax group (TrxG) of proteins. In contrast, Polycomb group (PcG) of proteins and heterochromatin factors maintain the transcriptionally repressed state of chromatin (facultative and constitutive heterochromatin, respectively). Together, they are critical for the establishment and maintenance of chromatin states throughout the development of eukaryotes (Ringrose and Paro 2004; Allshire and Madhani 2018; Schotta 2002). We examined the conservation of these factors in the genome of *L*. *boulardi*. Protein sequences from *D. melanogaster* were used as query sequences in a standalone BLAST to search against *L. boulardi* protein data set. The Polycomb group (PcG) and Trithorax group (TrxG) of proteins are well conserved in *L. boulardi* (Supplementary file 1: Table S8). However, unlike *Drosophila*, *polyhomeotic*, *extra sexcombs*, *pleiohomeotic* is present in only one copy. Heterochromatin factors, Heterochromatin protein 1 (HP1) family and Suppressor of variegation 3-9 (Su(var)3-9), the proteins that bind to and introduce heterochromatic histone methylation, respectively, are also conserved. Still, only one full length HP1 could be identified using blastp.

We further did a tblastn for identification of chromodomain and chromoshadow domain containing proteins in *L. boulardi*, since the characteristic feature of HP1 protein family is the presence of an N-terminal chromodomain and a C-terminal chromoshadow domain (Renato Paro 1990; R. Paro and Hogness 1991; Assland and Stewart 1995). All known chromodomain and chromoshadow domain sequences from *D. melanogaster* were used as seed sequences. A total of 49 proteins containing chromodomain were identified, which falls into four classes (Supplementary file 1: Table S9). All the chromodomain/chromoshadow domain sequences obtained were aligned with seed sequences from *D. melanogaster* using Clustal Omega (Madeira et al. 2019) followed by maximum likelihood tree generation with 1000 bootstrap steps using MEGA (Kumar et al. 2018) (Supplementary file 2: Figure S3 and S4). We identified only one HP1 protein (Class I) containing a chromodomain followed by a chromoshadow domain, eight such proteins but containing a single chromodomain (Class II). Four out of 49 have paired tandem chromodomain (Class IV) and 36 proteins contain chromodomain in combination with other domain families (Class III). A similar analysis done previously in ten hymenopterans has reported that Hymenopterans have a simple HP1 gene family comprising of one full HP1 and two partial HP1 genes (Fang, Schmitz, and Ferree 2015). However, we identified one full (chromodomain and chromoshadow domain) and eight partial (only chromodomain) HP1 homologs. The full HP1 protein identified is more similar to HP1b of *D. melanogaster* than to other paralogs. This indicates that the HP1 is more dynamic in *L. boulardi* than what is reported earlier in other hymenopterans.

## CONCLUSIONS

*Leptopilina* has been extensively used as a model system to study host-parasitoid biology. Our study presents a high-quality reference genome (375 Mb) of the specialist parasitoid wasp *Leptopilina boulardi* showing almost a complete coverage of the core gene repertoire shown by BUSCO analysis. A total of 25,259 protein-coding genes were predicted, out of which 22729 could be annotated using known protein signatures. We show that the genes responsible for determining the anteroposterior body axis (*Hox* genes) and sex determination are well conserved. *L. boulardi* has an incomplete DNA methylation toolkit; it is devoid of a *de novo* DNA methyltransferase (DNMT3). The HP1 family is much more diverse as compared to other hymenopterans. The other epigenetic regulators, Polycomb and trithorax group of proteins, are also retained. Overall, the basic machinery of epigenetic regulation is conserved, and though the unique features are noticed, their relevance needs further investigations.

The *L. boulardi* genome reported in this study provides a valuable resource to researchers studying parasitoids and can help shed light on the mechanisms of host-parasitoid interactions and understanding the immune response mechanisms in insects. The genome sequence of *L. boulardi* will also be a key element in understanding the evolution of parasitism in figitids. It will further enable genome editing and thereby advance the genetics of *L. boulardi*, facilitate the comparative studies of *Drosophila* parasitoids. More importantly, this resource fulfils the prerequisite for initiating research on epigenetic mechanisms underlying parasitism, and sex determination and other developmental mechanisms in *Leptopilina* genus.

## ABBREVIATIONS

PacBio: Pacific Biosciences
BUSCO: Benchmarking Universal Single-Copy Orthologs
RNA-Seq: RNA Sequencing
A-P: Anterior-Posterior
NCBI: National Center for Biotechnology Information
LD: Light/Dark
MaSuRCA: Maryland Super Read Celera Assembler
TRF: Tandem Repeats Finder
PERF: Perfect, Exhaustive Repeat Finder
SSR: Simple Sequence Repeat
STAR: Spliced Transcripts Alignment to a Reference
BLAST: Basic Local Alignment Search Tool
iTOL: Interactive Tree of Life
BACs: Bacterial Artificial Chromosomes
CDD: Conserved Domains Database
MEGA: Molecular Evolutionary Genetics Analysis

## AUTHORS’ CONTRIBUTIONS

R.K.M. conceived the study. R.K.M., S.K. and D.T.S. designed the project. S.K. prepared the samples and conducted experiments. D.T.S. performed genome assembly and annotation. D.T.S., S.K., A.S., and M.S. analyzed the data. S.K., D.T.S. wrote the manuscript. All authors read and approved the final manuscript.

## ACKNOWLEDGEMENTS

We thank Shubha Govind, The City College of the City University of New York, for providing us the Lb17 strain of *L. boulardi*. We acknowledge Indira Paddibhatla for introducing the *Drosophila – Leptopilina* system to our lab. Athira Rajeev is acknowledged for the initial quality control and processing of Illumina data.

